# RanGTP regulates the augmin complex

**DOI:** 10.1101/2022.12.23.521824

**Authors:** Jodi Kraus, Sophie M Travis, Matthew R King, Sabine Petry

## Abstract

Spindles are composed of microtubules that must nucleate at the right place and time during mitosis. Spindle microtubule nucleation is regulated by the GTPase Ran, which, through importin-αβ, releases a gradient of spindle assembly factors (SAFs) centered at chromosomes. Branching MT nucleation generates most spindle MTs and requires the augmin complex. In *Xenopus laevis*, Ran can control branching through the SAF TPX2, TPX2 is non-essential in other organisms. Thus, how Ran regulates branching MT nucleation in the absence of TPX2 is unknown. Here, we use in vitro pulldowns and TIRF microscopy to show that augmin is itself a SAF. Augmin directly interacts with both importins through two nuclear localization sequences on the Haus8 subunit, which overlap the MT binding site. Moreover, Ran controls localization of augmin to MTs in both *Xenopus* egg extract and in vitro. By uncovering that RanGTP directly regulates augmin, we demonstrate how Ran controls branching MT nucleation and, thereby, spindle assembly and cell division.

## INTRODUCTION

The mitotic spindle is a self-organized structure built from microtubules (MTs). Spindle MTs are made by several MT nucleation pathways that originate from various locations within the spindle (Meunier and Vernos, 2016; Petry, 2016; Wu and Akhmanova, 2017). MT nucleation in the cell requires the universal MT template, the γ-tubulin ring complex (γ-TuRC) (Flor-Parra et al., 2018; Gunzelmann et al., 2018; Oakley et al., 2015). However, different nucleation pathways use unique factors to recruit γ-TuRC to the appropriate MT nucleation site, and this recruitment is strictly regulated in space and time to allow proper spindle assembly (Sulimenko et al., 2022; Tovey and Conduit, 2018). In centrosomal spindles, chromosomes themselves are a key source of MTs, and, in acentrosomal spindles, chromosomes are the major regulator of spindle assembly (Cavazza and Vernos, 2016; Heald et al., 1996; Kalab and Heald, 2008). Central to the chromosome’s ability to make spindle MTs is a biochemical gradient of RanGTP (Caudron et al., 2005; Weaver and Walczak, 2015).

Ran is a soluble small guanosine triphosphatase (GTPase) of the Ras family (Cox and Der, 2010; Moore, 2001). During interphase, Ran directs the movement of cargoes, via effector heterodimeric importins, by promoting cargo uptake by importins in the cytoplasm and cargo release within the nucleus (Cavazza and Vernos, 2016; Dasso, 2001; Forbes et al., 2015). Once the nuclear envelope breaks down, the Ran-importin system regulates the formation of the mitotic spindle by directly controlling the release of spindle assembly factors (SAFs) (Cavazza and Vernos, 2016). In its active state, Ran is bound to GTP, which binds importin-β and causes importin-α to release NLS-containing cargoes, including SAFs (Cingolani et al., 1999; Lott and Cingolani, 2011). The inactive, GDP-bound Ran cannot bind the importin-αβ heterodimer and thus promotes cargo uptake and SAF sequestration (Nilsson et al., 2002; Ohba et al., 1999). Although Ran is distributed uniformly throughout the cell, RanGTP is concentrated at chromosomes, due to chromosomal localization of its activating guanosine nucleotide exchange factor, RCC1 (Carazo-Salas et al., 1999; Caudron et al., 2005). RanGTP diffuses away from chromosomes and encounters RanGTPase-activating proteins, allowing Ran to hydrolyze its bound GTP and become inactive (Caudron et al., 2005). The RanGTP gradient generates a secondary mitotic SAF gradient, where free active SAFs concentrate near chromosomes, thereby exerting spatial control of spindle assembly (Caudron et al., 2005; Tsuchiya et al., 2021; Weaver et al., 2015).

One key SAF that connects the RanGTP gradient to MT nucleation is the targeting protein for Xklp2 (TPX2). TPX2 facilitates branching MT nucleation, whereby one MT is nucleated from a pre-existing one, enabling amplification of MTs throughout the spindle (Petry and Vale, 2015; Travis et al., 2022b). In fact, branching provides a majority of MTs in centrosomal spindles (David et al., 2019) and is the main source of MTs in acentrosomal spindles including *Xenopus laevis* (Decker et al., 2018; Gouveia, 2022). Recent work showed that TPX2 forms a condensed phase that concentrates numerous branching factors at spindle MTs, as well as unpolymerized tubulin that can be used to build new MTs (King and Petry, 2020). Binding of TPX2 to the importin-αβ heterodimer inhibits both MT binding and condensation, thus inactivating TPX2 (Safari et al., 2021). However, recent evidence suggests that regulation of TPX2 may not be sufficient to control branching. In vitro studies of both *Drosophila* and human branching MT nucleation showed that branching can proceed in the absence of TPX2 (Tariq et al., 2020; Zhang et al., 2022), and in *Drosophila* cells TPX2 is not essential (Goshima, 2011). Thus, this begs the question of whether a second branching factor might be a Ran-regulated SAF.

The hetero-octameric augmin complex was first described in *Drosophila*, where subunits were shown to be required for robust spindle assembly (Goshima et al., 2008). Knockdown or depletion of augmin in vivo leads to dramatic reduction in spindle MT density, particularly in kinetochore fibers, and results in defects in spindle polarity and chromosome segregation both in *Drosophila* as well as in vertebrates and plants (David et al., 2019; Goshima et al., 2008; Ho et al., 2011; Lawo et al., 2009). Later, characterization of augmin in vitro demonstrated that augmin has binds to the universal MT nucleator, γ-TuRC, at one end of the complex known as tetramer III (T-III), while the other end of the complex, tetramer II (T-II) recognizes and binds MTs (Hsia et al., 2014; Song et al., 2018) (Figure 1a). Within T-II, there are two MT binding sites (Hsia et al., 2014; Travis et al., 2022a; Wu et al., 2008). The primary MT binding site was localized to the augmin subunit Haus8, within its intrinsically-disordered N-terminus (Hsia et al., 2014; Wu et al., 2008), while a second, minor MT binding site was very recently located within the Haus6 subunit (Travis et al., 2022a; Zupa et al., 2022). Thus, augmin promotes branching MT nucleation by recruiting γ-TuRC to the side of the MT (Alfaro-Aco et al., 2020; Song et al., 2018; Tariq et al., 2020; Zhang et al., 2022).

**Figure 1:**
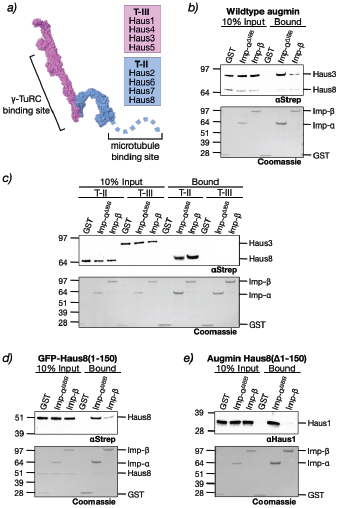
Augmin binds to importins. a) Structure of the augmin complex, broken into the γ-TuRC binding T-III (pink), comprised of subunits Haus1 and Haus3-5, and the MT-binding T-II (blue), comprised of subunits Haus2 and Haus6-8. The disordered N-terminus of Haus8 (shown as a dashed blue line) contains the primary MT binding site. Structure of *Xenopus* augmin was taken from [27]. b) Glutathione beads bound to either GST (control), GST-importin-α^ΔIBB^, or GST-importin-β were incubated with full-length augmin, then both the input and bound fraction were Western blotted for intact augmin complexes using an antibody against the Strep-tagged subunits Haus3 (T-III) and Haus8 (T-II). Below, GST and GST-importin loading was demonstrated by Coomassie stain. c) As in (b), importin-bound beads were incubated with augmin, either T-III or T-II, and binding of intact augmin subcomplex was detected via Western blot against the Strep-tagged subunits Haus3 and Haus8. d) Haus8^1-150^ (fused to an N-terminal Strep-tagged GFP) was incubated with importin-bound beads, and binding detected via Western blot. e) Augmin complex lacking the N-terminal 150 residues of Haus8 was incubated with importin-bound beads, and binding of intact augmin complex detected via Western blot against augmin subunit Haus1.

In this work, we demonstrate that the augmin complex, and more specifically the N-terminus of Haus8, have affinity for importin-αβ. This binding site, comprised of two conserved nuclear localization signal (NLS) sequences, overlaps with Haus8’s MT binding site, allowing importin to modulate augmin’s ability to bind to MTs. We further demonstrate that Ran is capable of releasing augmin from sequestration by importins. Thus, we identify augmin as a second point of Ran regulation within the branching MT nucleation pathway and, due to the fact that TPX2 is dispensable in many systems, propose that augmin may be the primary conserved point of Ran regulation in branching MT nucleation.

## RESULTS

### Augmin binds to importins via T-II and the N-terminus of Haus8

In order to determine whether augmin is a SAF, we first asked whether augmin is capable of binding importin-α and importin-β, a requirement of Ran regulation. Purified GST-tagged importins, as well as GST alone as a negative control, were immobilized on glutathione agarose resin and probed with recombinant *X. laevis* augmin complex. Because importin-α is autoinhibited by its NLS-like importin-β binding (IBB) sequence when not bound to importin-β (Lott and Cingolani, 2011), we used constitutively active importin-α lacking the IBB (residues 1-90), which we refer to as importin-α^ΔIBB^. We found that the augmin complex bound to both importin-α^ΔIBB^ and importin-β, with an apparent preference for importin-α^ΔIBB^ (Figure 1b).

Because the augmin complex is large (~450 kDa) and distinct functional roles are played by different subcomplexes (Song et al., 2018), we next asked where on the augmin complex the importin binding site or sites were located. We started by purifying the two separate soluble augmin subcomplexes, known as T-II and T-III (Figure 1a) (Song et al., 2018). We found that only T-II had affinity for the two importins (Figure 1c), binding both importin-α^ΔIBB^ and importin-β with approximately equal strength. In contrast, T-III displayed no appreciable binding to either importin (Figure 1c). Next, we analyzed the sequences of the four T-II subunits—Haus2, Haus6, Haus7, and Haus8—for potential importin binding sites (Kosugi et al., 2009). We found that the N-terminus of Haus8 (residues 1-150) has many of the unique hallmarks of importin binding regions, including predicted lack of structure and an abundance of basic amino acids. Fortuitously, as opposed to many other subunits of augmin, the N-terminus of Haus8 can be purified separately from the remainder of the augmin complex (Wu et al., 2008), and we found that this region does in fact bind both importin-α^ΔIBB^ and importin-β, again displaying a preference for importin-α^ΔIBB^ (Figure 1d).

After showing that Haus8^1-150^ was sufficient to bind to importins, we determined whether this was the sole importin binding site on augmin by expressing and purifying *Xenopus* augmin lacking Haus8^1-150^ and conducting the binding assay. Interestingly, we found that the truncated complex was still able to bind to importin-α^ΔIBB^, although importin-β binding was abrogated (Figure 1e). Based on our binding experiments with the two tetrameric subcomplexes, we suspect that this second binding site must be located somewhere within T-II, since importins do not bind T-III. However, we could not determine the location of this second importin-α binding region due to the lack of obvious predicted NLS sequences and because the entwined structure of the remaining T-II subunits prevents any other augmin fragments from folding correctly and thus retaining function (Gabel et al., 2022; Travis, 2022; Zupa, 2022).

### Haus8 contains two predicted NLS sequences that mediate importin-α binding

Having identified an importin binding region within Haus8, we sought to characterize exactly where those sites are located. Using an NLS prediction algorithm (Kosugi et al., 2009), we found two putative NLS sites in the N-terminus of Haus8, one comprising residues 23-54 and the other residues 138-148 (Figure 2a). The first putative NLS is bipartite and contains residues that are highly conserved among most vertebrates, whereas the second, monopartite NLS is predicted only in frogs and toads.

**Figure 2:**
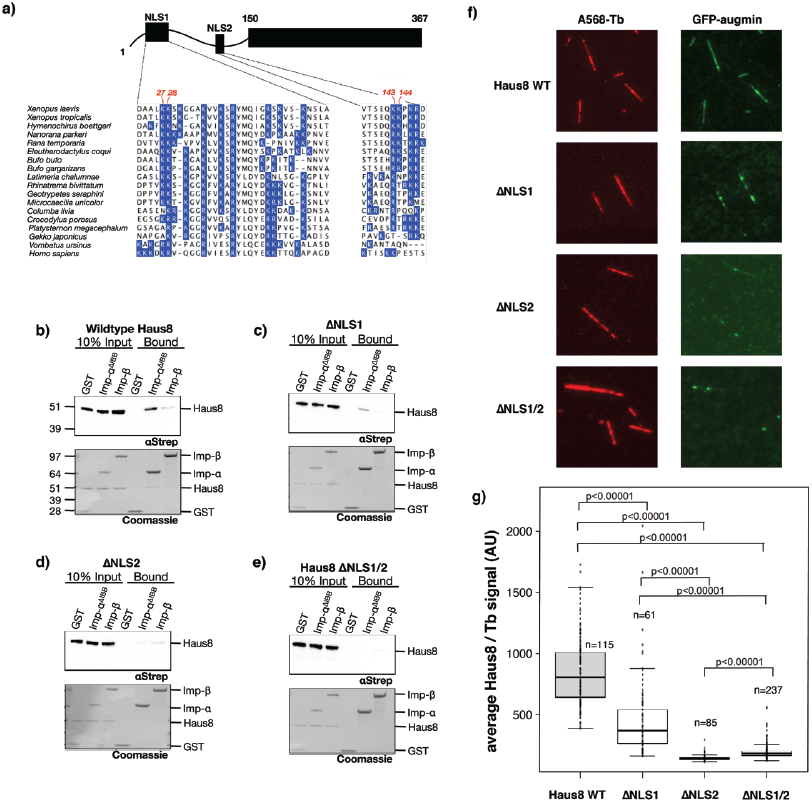
Haus8 of augmin binds to importins and MTs through two NLS sites. a) The augmin subunit Haus8 is predicted to contain two NLS sequences within its disordered N-terminus [25]. The domain architecture of Haus8 is cartooned at top, and the sequence of each predicted NLS in *Xenopus laevis* shown at bottom. *X. laevis* Haus8 is shown aligned to other vertebrate orthologues below, and all basic residues (arginine abbreviated as R, and lysine as K) are colored in blue. Indicated in red at the top of the sequence are the pairs of basic residues mutated to alanine to generate the Haus8 mutants ΔNLS1 (K27A/K28A) and ΔNLS2 (K143A/K144A). b)-e) Pulldowns of Strep-GFP-Haus8^1-150^, either wildtype or containing the indicated NLS mutants, were conducted as in Figure 1d. f) Selected TIRF images of in vitro binding of GST-GFP-Haus8^1-150^ to stabilized MT seeds. Haus8 constructs with mutated residues in NLS1 and/or NLS2 results in a reduction in binding, as quantified in g). g) Boxplot of average GFP-Haus8 signal relative to the average tubulin signal, where each marker represents a single MT from the experiment shown in f). The total number of MTs (n) was collected from two replicates. The boxes extend from the 25^th^ to 75^th^ percentile, and the upper and lower bars represent the minimum and maxiumum, respectively. P-values were calculated from independent T-tests.

To test whether one or both of these two putative NLS sites are required for importin binding, we mutated the NLSs by introducing double alanine mutations at either Lys-27 and Lys-28 or Lys-143 and Lys-144 (Makkerh et al., 1996; Zhang et al., 2011). We observed that mutation of either NLS led to a decrease in importin-α^ΔIBB^ binding relative to wildtype (Figure 2b). Moreover, mutation of NLS1 (Figure 2c) had a lesser impact on importin binding than mutation of NLS2 (Figure 2d). This may result either from a difference in importin-α binding affinity between the two NLS sequences, or, alternatively, may be because mutation of two residues in a bipartite NLS is less deleterious than mutation of two residues in a shorter, monopartite NLS. Nonetheless, combining mutations to both NLS sites completely abrogated importin-α^ΔIBB^ binding (Figure 2e). In contrast, mutation of either or both NLS sequences had little to no measurable effect on importin-β binding (Figure 2b-e). Thus, Haus8 contains two NLS regions recognized primarily by importin-α.

In the previously studied proteins TPX2 and NuMA, the NLS sites overlap with the MT binding domains (Chang et al., 2017; Haren and Merdes, 2002; Safari et al., 2021; Zhang et al., 2017). To test whether this is the case for augmin as well, we performed in vitro MT binding assays using biotinylated stabilized MT seeds attached via neutravidin to functionalized glass coverslips. GFP-labeled Haus8 NLS mutants were diluted to 100 nM, which mimics the endogenous concentration, and incubated with immobilized MT seeds. The degree of binding was assessed by total internal reflection fluorescence (TIRF) microscopy. We found that single NLS mutants decreased the binding to MT seeds, whereas the double mutant lacking both NLS sites inhibited binding altogether (Figure 2f-g). Thus, the MT binding sites in Haus8 overlap with the NLS regions.

### Importins prevent augmin from binding the MT lattice

A critical role of augmin is to bind to the side of the MT lattice in order to recruit γ-TuRC for branching MT nucleation. We hypothesized that this function may be Ran-regulated and therefore asked whether importin binding to augmin modulated its interactions with MTs. To address this, we performed in vitro MT binding assays with GFP-labeled Haus8. GFP-labeled Haus8 was incubated with a 10-fold molar excess of importins and then flowed onto the glass, followed by quantification of MT binding by TIRF microscopy. Interestingly, both importin-α and importin-β abrogated binding to MT seeds, indicating that importins indeed regulate augmin’s ability to bind to MTs (Figure 3a-b).

**Figure 3:**
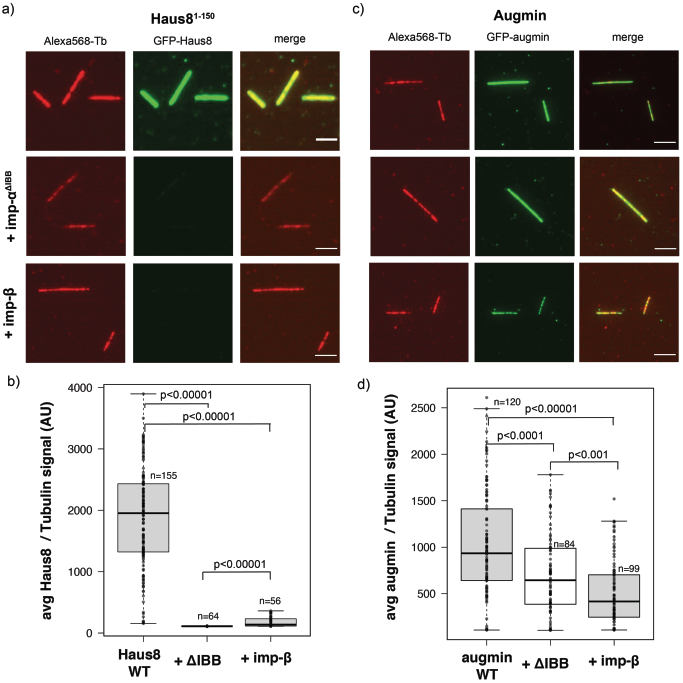
Importins regulate augmin binding to MTs. a) Wildtype (WT) GST-GFP-Haus8 localizes strongly to GMPCPP-stabilized MT seeds in vitro (top row), as visualized by TIRF microscopy. In the presence of importin-α^ΔIBB^ (middle row) or importin-β (bottom row), binding of Haus8 to MTs is diminished. This is quantified in b). b) Boxplot of average GFP-Haus8 signal relative to the average tubulin signal, where each marker represents a single MT from the experiment shown in f). The total number of MTs (n) was collected from two replicates. The boxes extend from the 25^th^ to 75^th^ percentile, and the upper and lower bars represent the minimum and maxiumum, respectively. P-values were calculated from independent T-tests. c) Wildtype GFP-labeled augmin localized to GMPCPP-stabilized MT seeds in vitro (top row), as visualized by TIRF microscopy. In the presence of importin-α^ΔIBB^ (middle row) or importin-β (bottom row), binding of augmin to MTs is decreased, but not eliminated. This is quantified in d). d) Boxplot of average GFP-augmin signal relative to the average tubulin signal, where each marker represents a single MT from the experiment shown in f). The total number of MTs (n) was collected from two replicates using two independent augmin preparations. The boxes extend from the 25^th^ to 75^th^ percentile, and the upper and lower bars represent the minimum and maxiumum, respectively. P-values were calculated from independent T-tests. Scale bars correspond to 5 μm.

We next tested how importins affected MT binding in the context of the full 8-subunit augmin complex to see if this behavior was reproduced. Interestingly, addition of both importin-α and importin-β significantly decreased binding of the full-length augmin to the MTs, but did not inhibit it altogether (Figure 3c-d). Therefore, this suggests that the secondary MT binding site in the Haus6 subunit is insensitive to importin binding, and still allows some localization to MTs. The functional consequences of the binding site in Haus6 is still under active investigation.

### Active Ran releases augmin from importins to promote MT binding in vitro and in *Xenopus* egg extract

SAF are not only characterized by importin inhibition, but also by the ability of RanGTP to release this inhibition, thereby activating the SAF. To assess whether Ran regulates MT binding of augmin, we tested whether augmin binds the importin-αβ heterodimer, which, in contrast to monomeric importins, is the predominant form importins take in the cell. In order to test this, we pulled down importin-αβ heterodimer using beads loaded with GFP-Haus8^1-150^. We found that the N-terminus of Haus8 bound strongly to a stoichiometric heterodimer of importin-α and importin-β relative to beads loaded only with GFP (Figure 4a). When to the mixture was added a 10-fold excess of Ran^Q69L^, a mutant of Ran locked in a GTP-bound conformational state, importin-αβ was lost from the resin, implying that Ran^Q69L^ triggers the release of augmin from importin-αβ (Figure 4a). Furthermore, importin-αβ inhibited binding of GFP-Haus8 to MT seeds in vitro under the same conditions, which was also rescued by the addition of active Ran^Q69L^ (Figure 4b-c).

**Figure 4:**
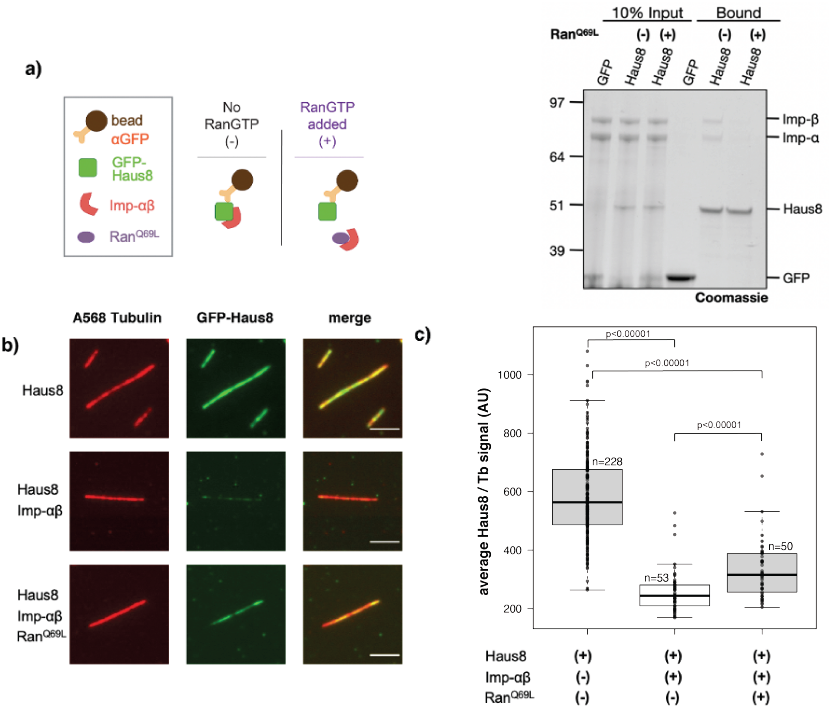
RanGTP releases importin inhibition of MT binding. a) GFP or GFP-Haus8^1-150^ was bound to α-GFP magnetic resin and incubated with importin-αβ in the presence or absence of a 10-fold excess of Ran^Q69L^. Both importin-αβ binding, as well as augmin loading, were assessed by Coomassie staining and the intensity of bands at the indicated sizes. b) In vitro localization of GST-GFP-Haus8^1-150^ binding to GMPCPP-stabilized MT seeds, as visualized by TIRF microscopy (top row). Addition of importin-αβ heterodimer inhibits binding of Haus8 to MTs (middle row), while addition of Ran^Q69L^ rescues MT binding of Haus8 (bottom row). This is quantified in c). c) Boxplot of average GFP-Haus8 signal relative to the average tubulin signal, where each marker represents a single MT from the experiment shown in f). n corresponds to the total number of MTs. The boxes extend from the 25^th^ to 75^th^ percentile, and the upper and lower bars represent the minimum and maxiumum, respectively. P-values were calculated from independent T-tests. Scale bars correspond to 5 μm.

To determine whether our results in vitro were representative of Ran regulation of augmin in a physiological environment, we next assessed the role of Ran in regulating augmin in *X. laevis* egg extract. To test this, we first took advantage of an antibody against Haus1 that pulls down the augmin complex without disrupting endogenous binding interactions with proteins including γ-TuRC (Song et al., 2018). Pulling down with this antibody, we found that substantially more importin-α and importin-β were bound to beads coated with α-Haus1 than non-specific IgG (Figure 5a), suggesting that augmin and importin-αβ interact in an endogenous setting. Additionally, binding of endogenous importin-αβ was regulated by Ran, as incubation of the beads with extract spiked with Ran^Q69L^ substantially reduced the amount of importin pulled down (Figure 5a). Finally, we generated branched MT structures in *X. laevis* egg extract and, using antibodies against Haus8 to visualize endogenous augmin, we found that augmin does not localize to MTs unless Ran^Q69L^ is present (Figure 5b).

**Figure 5:**
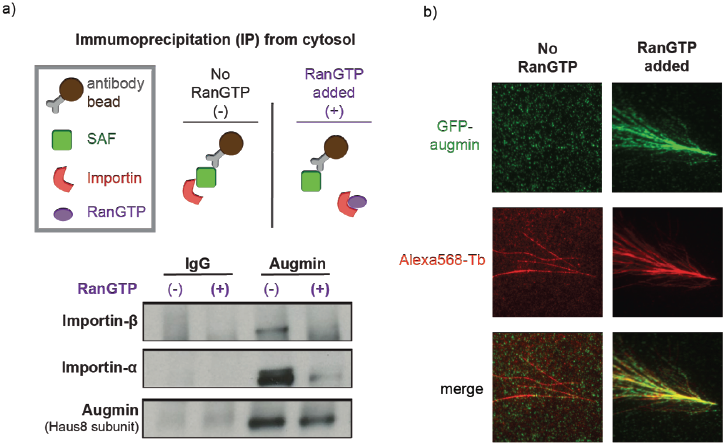
Ran regulates augmin in *Xenopus* egg extract. a) (top) Schematic of immunoprecipitation (IP) strategy where antibodies were conjugated to magnetic beads (top). (bottom) Western blot of IPs for a control antibody (IgG) and anti-augmin in the presence or absence of Ran^Q69L^. b) TIRF images of augmin in *Xenopus* egg extract in the absence (left column) and presence (right column) of Ran^Q69L^. Branching MT nucleation reactions were carried out with Alexa568-labeled tubulin and fixed after 15 minutes. Endogenous augmin was detected via indirect immunofluorescence with custom primary antibodies against Haus8 and Alexa488-conjugated secondary antibodies.

## DISCUSSION

Here we have shown that augmin is a bona fide SAF regulated by RanGTP. Augmin interacts with importin-α and importin-β, and independently binds each importin through two NLS sites, which we identified. The NLS sites are located in the disordered N-terminus of Haus8, which contains the primary MT binding site, indicating that the NLS sites and MT binding site(s) overlap. Moreover, importins inhibit binding of augmin to MTs, which can be reversed by addition of active RanGTP. Moreover, this regulatory system can be recapitulated both in vitro and in *Xenopus* egg extract.

While many studies have identified potential SAFs, none have previously recognized augmin as a candidate. High-throughput screens for importin clients in mammalian cells were limited to nuclear proteins (Yokoyama et al., 2008); therefore, unsurprisingly, augmin was not identified, as during interphase augmin localizes to the centrosome and is excluded from the nucleus (Lawo et al., 2009; Uehara et al., 2009). Other examples of SAFs that, during interphase, do not enter the nucleus include Kif2, which localizes to interphase MTs, and GM130, which localizes to the Golgi (Cavazza and Vernos, 2016). Permeability of the nuclear pore complex to importin-bound cargos decreases linearly with cargo mass, with the most efficient transport of proteins under 40 kDa (Lowe et al., 2015; Ma et al., 2012). This would explain why proteins like TPX2 can easily shuttle in and out of the interphase nucleus, but protein complexes like Kif2, GM130, and augmin cannot. Thus, many unidentified SAFs may exist and their identification may further explain how spindle assembly is regulated.

While our identification of augmin as a target of Ran regulation is new and unexpected, a rich literature exists to support Ran regulation of branching MT nucleation, through TPX2 (Giesecke and Stewart, 2010; Gruss et al., 2001; Safari et al., 2021; Schatz et al., 2003). Why might Ran separately regulate two branching factors? As discussed above, TPX2 is dispensable in certain systems, for example spindle assembly in *Drosophila*, which might be taken to suggest that, in these systems, Ran regulates branching exclusively through augmin. However, *Drosophila* augmin lacks the NLS-containing disordered N-terminus of its Haus8 subunit (known as Dgt4) and, in fact, previous work has suggested that, in *Drosophila*, augmin is not an importin client protein (Hayward et al., 2014). Thus, in these systems, it is more likely either that branching MT nucleation is independent of Ran or, as has been shown in *Drosophila*, is less reliant on branching MT nucleation for spindle assembly compared to e.g. human cells.

Conversely, in *Xenopus* and other species, both TPX2 and augmin are required for branching, and thus their regulation by Ran would seem to be redundant. Yet, true redundancy is a core organizational principle of spindle assembly, because it is so important for the cell’s survival. Redundancy can be seen from the single protein level, in the multiple NLSs found in augmin and TPX2 (Eibes et al., 2018), to the redundant functions of the various MT nucleation pathways. However, in addition, this dual control mechanism could be used to further sharpen the activity gradient of branching MT nucleation generated by RanGTP, and therefore provide more finely tuned spatiotemporal control of the pathway generating the majority of spindle MTs (ref).

In conclusion, our work, in identifying augmin as a Ran-regulated SAF, reveals another level of Ran control of branching MT nucleation, the major source of spindle MTs in vertebrate mitosis. However, it also raises larger questions in the regulation of spindle assembly as a whole, including how many unknown SAFs still are waiting to be identified and how simultaneous Ran-regulation of multiple SAFs shapes the landscape of MT nucleation during mitosis to, finally, generate the dynamic and complex structure of the mitotic spindle.

## ACKNOWLEDGEMENTS

We thank Mohammed Safari and Caitlin Lamb for providing reagents. We thank Venecia Valdez and Michael Rale for informative discussions and feedback. This work was supported by a fellowship from the Helen Hay Whitney Foundation (JK), National Institutes of Health grants F32GM142149 (SMT), NIH training grant 5T32GM007388-39 (MK), and R01 1R01GM141100-01A1 (SP).

## AUTHOR CONTRIBUTIONS

J.K. and S.M.T. conceived and performed experiments and wrote the manuscript. M.R.K. conceived and performed experiments. S.P. conceived experiments, provided expertise and feedback, and secured funding.

## METHODS

### Cloning

*Xenopus laevis* Haus8 codon-optimized for *E. coli* was synthesized by MacroLab and cloned into a pET3-based GFP-containing vector. N-terminal fragments were then subcloned via restriction enzyme cloning into modified pETDuet1 or pGEX4t1. NLS mutants were introduced through a modified QuickChange protocol (Liu and Naismith, 2008). Recombinant bacmids were created by transformation into DH10Bac competent cells (NEB), screening on XGAL plates, followed by bacmid isolation (Ciccarone, 1998) and confirmation through diagnostic PCR.

### Protein Expression and Purification

N-terminal Haus8 fragments, tagged either with Strep-GFP or GST-GFP, were expressed in Rosetta 2 (DE3) pLysS competent cells (NEB) grown in modified Terrific Broth at 27°C for 7 hours, following induction at mid-log with 0.5 mM isopropyl β-D-thiogalactopyranoside. After harvesting, cells were resuspended in lysis buffer (50 mM Tris pH 7.5, 750 mM NaCl, 5 mM ethylenediamine tetraacetic acid (EDTA), 1 mM dithiothreitol (DTT)) supplemented with 10 μg/mL DNase I from bovine pancreas (Roche) and 1 mM phenylmethylsulfonyl fluoride. Cells were lysed using an Avestin Emulsiflex C5, then clarified by ultracentrifugation at 105,000 x *g* at 4°C for 30 minutes. Clarified Strep-GFP-Haus8 lysate was batch loaded onto 2 mL StrepTactin Superflow resin (IBA) per liter of cells, washed with 50 column volumes of lysis buffer, and eluted with 5 column volumes of lysis buffer supplemented with 2.5 mM desthiobiotin. GST-GFP-Haus8 constructs were purified similarly, except using glutathione Superflow agarose resin (Pierce) and eluting with lysis buffer supplemented with 10 mM reduced glutathione instead of desthiobiotin. After elution, samples were concentrated using a 30 kDa molecular weight cutoff (MWCO) spin concentrator (Amicon) to 1 mL and loaded onto a Superdex 200 Increase 10/300 column (Cytiva) pre-equilibrated with high-salt CSFxB (10 mM HEPES pH 7.7, 500 mM KCl, 5 mM ethylene glycol-bis(β-aminoethyl ether)-N,N,N’,N’-tetraacetic acid, 1 mM MgCl_2_, 1 mM DTT). Fractions were pooled and concentrated prior to flash freezing in liquid nitrogen and storage at −80°C. Protein concentrations were determined by Bradford concentration assay using bovine serum albumin to generate a standard curve.

GST-tagged importins were previously described (King and Petry, 2020) and purified similarly, except substituting a lysis buffer with 200 mM NaCl and a gel filtration buffer with 100 mM KCl (low-salt CSFxB). Heterodimeric GST-importin-α bound to untagged importin-β was purified through separate expression and co-lysis of equal mass cell pellets of the two importins. Excess GST-importin-α was separated from stoichiometric heterodimer through gel filtration chromatography.

### His-human Ran^Q69L^ was purified as previously described (Alfaro-Aco et al., 2017)

Heterologous expression of *Xenopus laevis* augmin complex was performed in Sf9 cells as previously described (Song et al., 2018). Full-length augmin was expressed either with an N-terminal HRV3C-cleavable ZZ-tag on Haus6 and a C-terminal GFP-His tag on Haus2 as previously described, or with two additional Strep-GFP tags on both Haus3 and Haus8. T-II was expressed with an N-terminal ZZ-tag on Haus6, a C-terminal GFP-His tag on Haus2, and an N-terminal Strep-GFP tag on Haus8. T-III was expressed with an N-terminal Strep-GFP tag on Haus3. All complexes were purified in augmin lysis buffer (50 mM Tris pH 7.5, 200 mM NaCl, 5 mM EDTA, 2 μM β-mercaptoethanol, 10% (v/v) glycerol, 0.05% (v/v) Tween-20) supplemented with 10 μg/mL DNase I and 1 cOmplete protease inhibitor cocktail tablet (Roche) per L buffer. After lysis by Emulsiflex as above, lysates were clarified by ultracentrifugation at 250,000 x *g* at 4°C for 30 minutes. Clarified full-length and T-II lysates were batch loaded onto 1 mL IgG Sepharose resin, washed with 50 column volumes of lysis buffer, and eluted overnight by GST-HRV3C protease cleavage. T-III was purified similarly, except substituting StrepTactin resin and desthiobiotin elution as above. Following elution, complexes were either concentrated using a 50 kDa MWCO concentrator and further purified on a Superose 6 Increase 10/300 column (Cytiva) equilibrated in low-salt CSFxB or loaded directly onto Ni^2+^ agarose (Qiagen), eluted using 300 mM imidazole, then dialyzed overnight into low-salt CSFxB supplemented with 10% (w/v) D-sucrose.

### Pulldowns

For GST-importin-α^ΔIBB^ and GST-importin-β pulldowns, 30 μL of magnetic glutathione agarose (Pierce) was washed three times with 300 μL TBS supplemented with 0.1% (v/v) Tween-20 (TBS-T), then equilibrated into 300 μL low-salt CSFxB supplemented with 0.1% (v/v) Tween-20 (CSFxB-T). 30 μL of freshly made binding reaction, containing either 2 μM GST or GST-importin and either 2 μM GFP-Haus8 (1-150) or 200 nM augmin complex were added to the beads, allowed to mix, then incubated at 4°C for 2 hours. Supernatant was then removed and beads were washed three times with 300 μL CSFxB-T. Finally, beads were resuspended in 30 μL CSFxB-T, mixed with 10 μL 4x SDS-PAGE loading dye, and heated at 95°C to denature and elute any bound proteins. After SDS-PAGE, gels were either stained by Coomassie to detect total protein content, or transferred to nitrocellulose via iBLOT (Invitrogen) and probed for augmin using either murine α-Strep (Qiagen) or a previously published rabbit α-*Xenopus* Haus1 (Song et al., 2018).

For pulldowns with GST-importin-α bound to untagged importin-β, 30 μL of GFPTrap magnetic agarose (Chromotek) was substituted for glutathione resin above, and TBS-T and CSFxB-T were supplemented with 10 mg/mL κ-casein; however, the remainder of the pulldown proceeded in the same manner.

For pulldowns of endogenous augmin, *Xenopus* egg extract was prepared as previously described (Alfaro-Aco et al., 2017; Song et al., 2018). α-Haus8 [21] was immobilized on magnetic protein A Dynabeads (Thermo Fisher Scientific), then pulldowns were carried out according to previously described protocols (Alfaro-Aco et al., 2017). In addition to probing with α-Haus8, pulldown blots were probed using custom antibodies against *X. laevis* GST-importin-α and GST-importin-β generated by GenScript and purified from serum according to previously published protocols (Alfaro-Aco et al., 2017; Song et al., 2018).

### Tubulin Labeling and Polymerization of GMPCPP-Stabilized Microtubules

Bovine brain tubulin was labeled following prior methods (Hyman et al., 1991). Using Alexa568-NHS ester (Invitrogen, A20003) yielded 36-40% labeling efficiency. Single cycled GMPCPP-stabilized MTs were made as previously described (Alfaro-Aco et al., 2020; Gell et al., 2010). Briefly, 12 μM unlabeled bovine tubulin supplemented with 1 μM Alexa-568 tubulin and 1 μM biotin tubulin was polymerized in BRB80 buffer (80 mM PIPES, 1 mM EGTA, 1 mM MgCl_2_) in the presence of 1 mM GMPCPP for 1 hour at 37°C. After 1 hour, the MT seed mixture was centrifuged at 13,000 x *g* for 15 minutes. The supernatant was removed and the pellet was resuspended in warm BRB80 buffer supplemented with 1 mM GMPCPP.

### Preparation of Polyethylene Glycol (PEG)-Functionalized Coverslips

22 mm x 22 mm cover glasses (Carl Zeiss, 474030-9020-000) were silanized and reacted with PEG as previously described (Bieling et al., 2010), except that hydroxyl-PEG-3000-amine and biotin-PEG-3000-amine were used. Glass slides were passivated using poly(L-lysine)-PEG. Flow chambers for TIRF microscopy were prepared using parafilm and gentle heating to seal coverslips to the glass slides.

### Attachment of GMPCPP-stabilized Microtubules to PEG-Functionalized Coverslips

Flow chambers were incubated with 5% Pluronic F-127 in water (Invitrogen, P6866) for 5 minutes at room temperature and then washed with assay buffer (BRB80, 5 mM β-mercaptoethanol, 0.075% (w/v) methylcellulose, 1% (w/v) glucose, 0.02% (v/v) Brij-35 (Thermo Scientific, 20150)) supplemented with 50 μg/mL κ-casein. Flow chambers were then incubated with an assay buffer containing 50 μg/mL NeutrAvidin (Invitrogen, A2666) for 2 minutes on a metal block on ice and subsequently washed with BRB80. Next, flow chambers were incubated for 5 minutes at room temperature with Alexa-568 labeled, biotinylated GMPCPP-stabilized MTs diluted 1:2000 in BRB80. Unbound MTs were removed by additional BRB80 washes.

### Binding of Proteins to GMPCPP-Stabilized MTs

To assess the binding of augmin and Haus8 constructs to MTs, augmin and Haus8 were diluted to 100 nM in CSFxB with 100 mM KCl and added to the flow chamber containing GMPCPP-stabilized microtubule seeds, previously attached to the coverslip surface. This was incubated for 10 minutes at room temperature. Unbound proteins were washed away using BRB80. For experiments with importins, importin proteins were incubated with augmin proteins on ice for 10 minutes prior to entering the flow chamber. All samples were imaged immediately.

### TIRF Microscopy and Image Analysis

Total internal reflectance fluorescence (TIRF) microscopy was performed with a Nikon TiE microscope using a 100 × 1.49 NA objective. Andor Zyla cCMOS camera was used for acquisition, with a field of view 165.1 × 139.3 μM. Multi-color images were collected using the NIS-Elements software (Nikon). All adjustable parameters for imaging (exposure time, laser intensity, and TIRF angle) were kept the same within experiments. For in vitro experiments, the objective was warmed with an objective heater at 33°C. Images belonging to the same experiment were contrast-matched. Images used for quantification of MT binding were analyzed using ImageJ. To segment microtubules, the tubulin signal was thresholded via the Otsu method. MTs were isolated from the mask by setting the minimum particle area as 0.5 μm^2^. Average fluorescent signal per pixel was recorded for each MT with and without additional proteins. To compare augmin fluorescence intensity across experiments, the intensity was normalized with respect to the tubulin signal.

### *Xenopus* egg extract reactions

Branching MT nucleation reactions were carried out in 5 μL volume flow cells with Alexa568-labeled fluorescent tubulin in the presence or absence of Ran^Q69L^, as previously described (Petry et al., 2013). After 15 minutes, fixative (−20°C methanol) was added and allowed to incubate for 1 minute. This lead to branched MTs adhered to the coverslip within each flow cell (hereafter called reactions). Fixative was washed out with a continuous flow of 50 μl blocking buffer (5% normal goat serum S1000 in low-salt CSFxB; Vector Labs), then reactions were incubated in that buffer for 1 hour at 4°C. After this time, reactions were incubated overnight at 4°C in blocking buffer with custom rabbit polyclonal α-Haus8. The following day, reactions were subjected to three rounds of washing with 50 μl low-salt CSFxB each followed by a 15 minute incubation. Reactions were incubated for 1 hour in blocking buffer containing goat αrabbit IgG (H+L) secondary antibody Alexa Fluor 568 conjugate (Thermo Fisher Scientific). Again, reactions were washed with 50 μl low-salt CSFxB three times, and each wash was followed by a 15 minute incubation. Finally, reactions were mounted with ProLong Diamond Antifade Mountant (Life Technologies) which was allowed to cure before imaging (typically >2 hours). All steps and buffers were carried out at room temperature unless otherwise specified, and all incubations were performed in a humidity chamber.

## REFERENCES

Alfaro-Aco, R., A. Thawani, and S. Petry. 2017. Structural analysis of the role of TPX2 in branching microtubule nucleation. J. Cell Biol. 216:983–997.

Alfaro-Aco, R., A. Thawani, and S. Petry. 2020. Biochemical reconstitution of branching microtubule nucleation. Elife. 9.

Bieling, P., I.A. Telley, C. Hentrich, J. Piehler, and T. Surrey. 2010. Fluorescence microscopy assays on chemically functionalized surfaces for quantitative imaging of microtubule, motor, and +TIP dynamics. Methods Cell Biol. 95:555–580.

Carazo-Salas, R.E., G. Guarguaglini, O.J. Gruss, A. Segref, E. Karsenti, and I.W. Mattaj. 1999. Generation of GTP-bound Ran by RCC1 is required for chromatin-induced mitotic spindle formation. Nature. 400:178–181.

Caudron, M., G. Bunt, P. Bastiaens, and E. Karsenti. 2005. Spatial coordination of spindle assembly by chromosome-mediated signaling gradients. Science. 309:1373–1376.

Cavazza, T., and I. Vernos. 2016. The RanGTP Pathway: From Nucleo-Cytoplasmic Transport to Spindle Assembly and Beyond. Front. Cell Dev. Biol. 3.

Chang, C.C., T.L. Huang, Y. Shimamoto, S.Y. Tsai, and K.C. Hsia. 2017. Regulation of mitotic spindle assembly factor NuMA by Importin-beta. J. Cell Biol. 216:3453–3462.

Ciccarone, V.C., Polayes, D.A., Luckow, V.A. 1998. Generation of recombinant baculovirus DNA in E. coli using a baculovirus shuttle vector. Methods Mol Med. 13:213–235.

Cingolani, G., C. Petosa, K. Weis, and C.W. Muller. 1999. Structure of importin-beta bound to the IBB domain of importin-alpha. Nature. 399:221–229.

Cox, A.D., and C.J. Der. 2010. Ras history: The saga continues. Small GTPases. 1:2–27.

Dasso, M. 2001. Running on ran: Nuclear transport and the mitotic spindle. Cell. 104:321–324.

David, A.F., P. Roudot, W.R. Legant, E. Betzig, G. Danuser, and D.W. Gerlich. 2019. Augmin accumulation on long-lived microtubules drives amplification and kinetochore-directed growth. J. Cell Biol. 218:2150–2168.

Decker, F., D. Oriola, B. Dalton, and J. Brugues. 2018. Autocatalytic microtubule nucleation determines the size and mass of Xenopus laevis egg extract spindles. Elife. 7.

Eibes, S., N. Gallisa-Sune, M. Rosas-Salvans, P. Martinez-Delgado, I. Vernos, and J. Roig. 2018. Nek9 phosphorylation defines a new role for TPX2 in Eg5-dependent centrosome separation before nuclear envelope breakdown. Curr. Biol. 28:121.

Flor-Parra, I., A.B. Iglesias-Romero, and F. Chang. 2018. The XMAP215 ortholog Alp14 promotes microtubule nucleation in fission yeast. Current Biology. 28:1681–1691.

Forbes, D.J., A. Travesa, M.S. Nord, and C. Bernis. 2015. Nuclear transport factors: global regulation of mitosis. Curr. Op. Cell Biol. 35:78–90.

Gabel, C.A., Z. Li, A.G. DeMarco, Z.G. Zhang, J. Yang, M.C. Hall, D. Barford, and L.F. Chang. 2022. Molecular architecture of the augmin complex. Nat. Commun. 13.

Gell, C., V. Bormuth, G.J. Brouhard, D.N. Cohen, S. Diez, C.T. Friel, J. Helenius, B. Nitzsche, H. Petzold, J. Ribbe, E. Schaffer, J.H. Stear, A. Trushko, V. Varga, P.O. Widlund, M. Zanic, and J. Howard. 2010. Microtubule dynamics reconstituted in vitro and imaged by single-molecule fluorescence microscopy. Methods Cell Biol. 95:221–245.

Giesecke, A., and M. Stewart. 2010. Novel binding of the mitotic regulator TPX2 (Target Protein for Xenopus Kinesin-like Protein 2) to importin-alpha. J. Biol. Chem. 285:17628–17635.

Goshima, G. 2011. Identification of a TPX2-Like Microtubule-Associated Protein in Drosophila. Plos One. 6.

Goshima, G., M. Mayer, N. Zhang, N. Stuurman, and R.D. Vale. 2008. Augmin: a protein complex required for centrosome-independent microtubule generation within the spindle. J. Cell Biol. 181:421–429.

Gouveia, B., Setru, S.U., King, M.R., Stone, H.A., Shaevitz, J.W., Petry, S. 2022. Acentrosomal spindles assemble from branching microtubule nucleation near chromosomes Biorxiv.

Gruss, O.J., R.E. Carazo-Salas, C.A. Schatz, G. Guarguaglini, J. Kast, M. Wilm, N. Le Bot, I. Vernos, E. Karsenti, and I.W. Mattaj. 2001. Ran induces spindle assembly by reversing the inhibitory effect of importin alpha on TPX2 activity. Cell. 104:83–93.

Gunzelmann, J., D. Ruthnick, T.C. Lin, W.L. Zhang, A. Neuner, U. Jakle, and E. Schiebel. 2018. The microtubule polymerase Stu2 promotes oligomerization of the gamma-TuSC for cytoplasmic microtubule nucleation. Elife. 7.

Haren, L., and A. Merdes. 2002. Direct binding of NuMA to tubulin is mediated by a novel sequence motif in the tail domain that bundles and stabilizes microtubules. J. Cell Sci. 115:1815–1824.

Hayward, D., J. Metz, C. Pellacani, and J.G. Wakefield. 2014. Synergy between multiple microtubule-generating pathways confers robustness to centrosome-driven mitotic spindle formation. Dev. Cell. 28:81–93.

Heald, R., R. Tournebize, T. Blank, R. Sandaltzopoulos, P. Becker, A. Hyman, and E. Karsenti. 1996. Self-organization of microtubules into bipolar spindles around artificial chromosomes in Xenopus egg extracts. Nature. 382:420–425.

Ho, C.M.K., T. Hotta, Z.S. Kong, C.J.T. Zeng, J. Sun, Y.R.J. Lee, and B. Liu. 2011. Augmin plays a critical role in organizing the spindle and phragmoplast microtubule arrays in Arabidopsis. Plant Cell. 23:2606–2618.

Hsia, K.C., E.M. Wilson-Kubalek, A. Dottore, Q. Hao, K.L. Tsai, S. Forth, Y. Shimamoto, R.A. Milligan, and T.M. Kapoor. 2014. Reconstitution of the augmin complex provides insights into its architecture and function. Nat. Cell Biol. 16:852–863.

Hyman, A., D. Drechsel, D. Kellogg, S. Salser, K. Sawin, P. Steffen, L. Wordeman, and T. Mitchison. 1991. Preparation of modified tubules. Methods in Enzymology. 196:478–485.

Kalab, P., and R. Heald. 2008. The RanGTP gradient - a GPS for the mitotic spindle. J. Cell Sci. 121:1577–1586.

King, M.R., and S. Petry. 2020. Phase separation of TPX2 enhances and spatially coordinates microtubule nucleation. Nat. Commun. 11.

Kosugi, S., M. Hasebe, M. Tomita, and H. Yanagawa. 2009. Systematic identification of cell cycle-dependent yeast nucleocytoplasmic shuttling proteins by prediction of composite motifs. Proc. Natl. Acad. Sci. U.S.A. 106:10171–10176.

Lawo, S., M. Bashkurov, M. Mullin, M.G. Ferreria, R. Kittler, B. Habermann, A. Tagliaferro, I. Poser, J.R.A. Hutchins, B. Hegemann, D. Pinchev, F. BuchholZ, J.M. Peters, A.A. Hyman, A.C. Gingras, and L. Pelletier. 2009. HAUS, the 8-subunit human augmin complex, regulates centrosome and spindle integrity. Curr. Biol. 19:816–826.

Liu, H.T., and J.H. Naismith. 2008. An efficient one-step site-directed deletion, insertion, single and multiple-site plasmid mutagenesis protocol. Bmc Biotech. 8.

Lott, K., and G. Cingolani. 2011. The importin beta binding domain as a master regulator of nucleocytoplasmic transport. Biochim Biophys Acta. 1813:1578–1592.

Lowe, A.R., J.H. Tang, J. Yassif, M. Graf, W.Y.C. Huang, J.T. Groves, K. Weis, and J.T. Liphardt. 2015. Importin-beta modulates the permeability of the nuclear pore complex in a Ran-dependent manner. Elife. 4.

Ma, J., A. Goryaynov, A. Sarma, and W.D. Yang. 2012. Self-regulated viscous channel in the nuclear pore complex. Proc. Natl. Acad. Sci. U.S.A. 109:7326–7331.

Makkerh, J.P.S., C. Dingwall, and R.A. Laskey. 1996. Comparative mutagenesis of nuclear localization signals reveals the importance of neutral and acidic amino acids. Curr. Biol. 6:1025–1027.

Meunier, S., and I. Vernos. 2016. Acentrosomal Microtubule Assembly in Mitosis: The Where, When, and How. Trends Cell Biol. 26:80–87.

Moore, J.D. 2001. The Ran-GTPase and cell-cycle control. Bioessays. 23:77–85.

Nilsson, J., K. Weis, and J. Kjems. 2002. The C-terminal extension of the small GTPase Ran is essential for defining the GDP-bound form. J Mol Biol. 318:583–593.

Oakley, B.R., V. Paolillo, and Y.X. Zheng. 2015. gamma-Tubulin complexes in microtubule nucleation and beyond. Mol. Biol. Cell. 26:2957–2962.

Ohba, T., M. Nakamura, H. Nishitani, and T. Nishimoto. 1999. Self-organization of microtubule asters induced in Xenopus egg extracts by GTP-bound Ran. Science. 284:1356–1358.

Petry, S. 2016. Mechanisms of Mitotic Spindle Assembly. Annu Rev Biochem. 85:659–683.

Petry, S., A.C. Groen, K. Ishihara, T.J. Mitchison, and R.D. Vale. 2013. Branching microtubule nucleation in Xenopus egg extracts mediated by augmin and TPX2. Cell. 152:768–777.

Petry, S., and R.D. Vale. 2015. Microtubule nucleation at the centrosome and beyond. Nat Cell Biol. 17:1089–1093.

Safari, M.S., M.R. King, C.P. Brangwynne, and S. Petry. 2021. Interaction of spindle assembly factor TPX2 with importins-alpha/beta inhibits protein phase separation. J. Biol. Chem. 297.

Schatz, C.A., R. Santarella, A. Hoenger, E. Karsenti, I.W. Mattaj, O.J. Gruss, and R.E. Carazo-Salas. 2003. Importin alpha-regulated nucleation of microtubules by TPX2. Embo J. 22:2060–2070.

Song, J.G., M.R. King, R. Zhang, R.S. Kadzik, A. Thawani, and S. Petry. 2018. Mechanism of how augmin directly targets the gamma-tubulin ring complex to microtubules. J. Cell Biol. 217:2417–2428.

Sulimenko, V., E. Draberova, and P. Draber. 2022. gamma-Tubulin in microtubule nucleation and beyond. Front Cell Dev Biol. 10:880761.

Tariq, A., L. Green, J.C.G. Jeynes, C. Soeller, and J.G. Wakefield. 2020. In vitro reconstitution of branching microtubule nucleation. Elife. 9.

Tovey, C.A., and P.T. Conduit. 2018. Microtubule nucleation by gamma-tubulin complexes and beyond. Essays Biochem. 62:765–780.

Travis, S.M., B.P. Mahon, W. Huang, M. Ma, M.J. Rale, J. Kraus, D.J. Taylor, R. Zhang, and S. Petry. 2022a. Integrated Model of the Vertebrate Augmin Complex. Preprint at bioRxiv https://doi.org/10.1101/2022.09.26.509603.

Travis, S.M., B.P. Mahon, and S. Petry. 2022b. How Microtubules Build the Spindle Branch by Branch. Annu Rev Cell Dev Biol. 38:1–23.

Travis, S.M., Mahon, B.P., Huang, W., Ma, M., Rale, M.J., Kraus, J., Taylor, D.J., Zhang, R., Petry, S. 2022. Integrated model of the vertebrate augmin complex. BioRxiv.

Tsuchiya, K., H. Hayashi, M. Nishina, M. Okumura, Y. Sato, M.T. Kanemaki, G. Goshima, and T. Kiyomitsu. 2021. Ran-GTP Is Non-essential to Activate NuMA for Mitotic Spindle-Pole Focusing but Dynamically Polarizes HURP Near Chromosomes. Curr Biol. 31:115–127 e113.

Uehara, R., R.S. Nozawa, A. Tomioka, S. Petry, R.D. Vale, C. Obuse, and G. Goshima. 2009. The augmin complex plays a critical role in spindle microtubule generation for mitotic progression and cytokinesis in human cells. Proc. Natl. Acad. Sci. U.S.A. 106:6998–7003.

Weaver, L.N., S.C. Ems-McClung, S.H. Chen, G. Yang, S.L. Shaw, and C.E. Walczak. 2015. The Ran-GTP gradient spatially regulates XCTK2 in the spindle. Curr Biol. 25:1509–1514.

Weaver, L.N., and C.E. Walczak. 2015. Spatial gradients controlling spindle assembly. Biochem Soc Trans. 43:7–12.

Wu, G.K., Y.T. Lin, R. Wei, Y.M. Chen, Z.Y. Shan, and W.H. Lee. 2008. Hice1, a novel microtubule-associated protein required for maintenance of spindle integrity and chromosomal stability in human cells. Mol. Cell Biol. 28:3652–3662.

Wu, J., and A. Akhmanova. 2017. Microtubule-Organizing Centers. Annu Rev Cell Dev Biol. 33:51–75.

Yokoyama, H., O.J. Gruss, S. Rybina, M. Caudron, M. Schelder, M. Wilm, I.W. Mattaj, and E. Karsenti. 2008. Cdk11 is a RanGTP-dependent microtubule stabilization factor that regulates spindle assembly rate. J Cell Biol. 180:867–875.

Zhang, R., J. Roostalu, T. Surrey, and E. Nogales. 2017. Structural insight into TPX2-stimulated microtubule assembly. Elife. 6.

Zhang, Y., X. Hong, S. Hua, and K. Jiang. 2022. Reconstitution and mechanistic dissection of the human microtubule branching machinery. J. Cell Biol. 7:e202109053.

Zhang, Z.C., N. Satterly, B.M.A. Fontoura, and Y.M. Chook. 2011. Evolutionary development of redundant nuclear localization signals in the mRNA export factor NXF1. Mol. Biol. Cell. 22:4657–4668.

Zupa, E., M. Wurtz, A. Neuner, T. Hoffmann, M. Rettel, A. Bohler, B.J.A. Vermeulen, S. Eustermann, E. Schiebel, and S. Pfeffer. 2022. The augmin complex architecture reveals structural insights into microtubule branching. Nat Commun. 13:5635.

Zupa, E., Würtz, M., Neuner, A., Hoffmann, T., Rettel, M., Böhler, A., Vermeulen, B.J.A., Eustermann, S., Schiebel, E., Pfeffer, S. 2022. The augmin complex architecture reveals structural insights into microtubule branching. Nat. Commun. 13.

